# Widespread innervation of motoneurons by spinal V3 neurons globally amplifies locomotor output in mice

**DOI:** 10.1101/2024.03.15.585199

**Authors:** Han Zhang, Dylan Deska-Gauthier, Colin S. MacKay, Krishnapriya Hari, Ana M. Lucas-Osma, Joanna Borowska-Fielding, Reese L. Letawsky, Turgay Akay, Keith K. Fenrich, David J. Bennett, Ying Zhang

## Abstract

While considerable progress has been made in understanding the neuronal circuits that underlie the patterning of locomotor behaviours such as walking, less is known about the circuits that amplify motoneuron output to enable adaptable increases in muscle force across different locomotor intensities. Here, we demonstrate that an excitatory propriospinal neuron population (V3 neurons, Sim1^+^) forms a large part of the total excitatory interneuron input to motoneurons (∼20%) across all hindlimb muscles. Additionally, V3 neurons make extensive connections among themselves and with other excitatory premotor neurons (such as V2a neurons). These circuits allow local activation of V3 neurons at just one segment (via optogenetics) to rapidly depolarize and amplify locomotor-related motoneuron output at all lumbar segments in both the in vitro spinal cord and the awake adult mouse. Interestingly, despite similar innervation from V3 neurons to flexor and extensor motoneuron pools, functionally, V3 neurons exhibit a pronounced bias towards activating extensor muscles. Furthermore, the V3 neurons appear essential to extensor activity during locomotion because genetically silencing them leads to slower and weaker mice with a poor ability to increase force with locomotor intensity, without much change in the timing of locomotion. Overall, V3 neurons increase the excitability of motoneurons and premotor neurons, thereby serving as global command neurons that amplify the locomotion intensity.

## Introduction

The generation of coordinated movement involves an intricate interplay between the brain, spinal cord, muscles and limbs, allowing animals to perform a wide range of movements – from reflexive responses to complex voluntary actions^1,2^. At the heart of this system is the spinal cord, a critical hub for transmitting neural signals between the brain and body, orchestrating a wide array of animal behaviours, including locomotion^3–6^. The complexity of the spinal neuronal circuits controlling locomotor output is epitomized by the highly diverse spinal interneurons and motoneurons that form the spinal central pattern generator (CPG) for rhythmic and patterned locomotion^7,8^. Multiple molecularly identified cardinal interneuron classes play distinctive roles in the regulation of locomotor behaviours, such as those contributing to the rhythm generation, and others ensuring the correct timing of the activation patterns of different muscles, including left-right and flexion-extension alternation^4,7,8^. While much is known about how locomotor rhythms and patterns are produced by the CPG, less is known about how the output of the CPG is regulated at the motoneuron level to ultimately amplify motoneuron output and allow locomotor activities of differing intensities to be reliably produced.

There are several genetically identifiable classes of spinal interneurons that have direct projections to motoneurons (premotor neurons), enabling them to modulate motor output, including V0, V1, V2, and V3 neurons^9–12^. However, many of these premotor neurons have net inhibitory effects on motoneurons and so cannot directly amplify motoneuron output. Moreover, most of them are crucial in the regulation of detailed timing/patterning of the CPG output through inhibiting unwanted motor activity. For instance, the V1 and V2b neurons are strictly inhibitory and responsible for reciprocal flexor-extensor alternation^13–18^, while V0d and V0v neurons ensure the left-right alternation of hindlimbs at different speeds via direct or indirect inhibition to motoneurons^19–23^. Even V2a neurons, an excitatory and ipsilaterally projecting population, have only been shown to be responsible for hindlimb left-right alternation without affecting locomotor intensity^24–28^. While one subpopulation of V0 neurons, V0c, has been shown to regulate the intensity of locomotion (during swimming), it does so indirectly by its unique cholinergic C boutons that modulate intrinsic motoneuron properties via the G protein-coupled muscarinic receptor^29^. Overall, the V0d, V0v, V1, V2b and V2a neurons are thus crucial for the detailed timing of rhythmic CPG locomotor activity, with all but V2a contributing inhibition.

Here, we suggest that one major function of V3 neurons is quite different from that of most other premotor neurons, not so much involved in regulating the detailed timing of rhythmic activity, but instead broadly amplifying the output of many motoneuron types, as we explore in this paper. V3 neurons are glutamatergic excitatory neurons that are mostly contralaterally projecting and characterized by the embryonic expression of the transcription factor Sim1^30^. V3 neurons are highly heterogeneous, with molecularly, anatomically and electrophysiologically distinct properties, ultimately forming functionally distinct subpopulations with dorsal, intermediate and ventral locations and differing molecular identities^31–34^. Although we have recently begun to study the role of the more dorsal V3 neurons with ascending projections^35^, we know less about the function of the ventral V3 neurons (V3v), which make up the majority of the total V3 population^34^. V3v neurons are located in lamina VIII and X, and mostly form long commissural axonal projections that travel across the entire contralateral lumbosacral spinal cord. They have both descending and ascending projections, though the descending projections dominate^32–34^. While most V3v neurons project to contralateral segments, a small subset of V3v neurons form ipsilateral projections to motoneurons, though these are strictly local and represent only a small percent of total motoneuron innervation (ipsilateral)^34,36^. Given the anatomical similarity of V3_V_ neurons to commissural laminae VIII neurons that innervate contralateral motoneurons, as defined by Jankowska decades ago^37^, these two populations are likely the same neurons. Accordingly, V3v neurons likely form the bulk of the excitatory input to contralateral motoneurons and receive extensive descending reticulospinal (RST), vestibulospinal (VST) and sensory inputs, as shown for commissural lamina VIII neurons^38^. V3v neurons may, therefore, serve as an important target for supraspinal amplification of motoneuron output, suggesting a functional role unique from all other previously described ventral premotor neuron classes.

Indeed, genetic silencing of V3 neurons does not much affect left-right or flexor-extensor alternation, but leads to weaker rhythmic locomotor activity that is harder to initiate in the isolated neonatal spinal cords^30^ and slower unstable gaits in walking mice^35^. On the other hand, optogenetic activation of V3 neurons in neonatal spinal cord preparations broadly increases the contralateral extensor motoneuron activity during drug-induced locomotor-related activity^39^. Interestingly, a recent study in zebrafish using activity-dependent imaging demonstrates that V3 neurons do not play a role in rhythmic swimming generation, but instead become tonically active during bouts of swimming, boosting overall motor activity, presumably bringing motoneurons closer to threshold to amplify rhythmic CPG output^40^. Taken together, these findings suggest that V3 neurons may play a key role in controlling the gain of motoneuron output, globally depolarizing motoneurons and premotor neurons. This should ultimately directly activate motoneurons and indirectly amplify the action of the CPG by bringing motoneurons closer to threshold.

To explore this gain control hypothesis, here we systematically examined the anatomical and functional relationships between V3 neurons and motoneurons, both *in vitro* and *in vivo*. By employing tract tracing, genetic labelling, and activating and silencing of V3 neurons, we demonstrate, as expected, that V3 neurons help amplify motor output during locomotion. However, we were surprised that V3 neurons produce a remarkably large portion of the total input to all hindlimb motoneurons, enabling them to globally and synchronously increase motoneuron activity, regardless of the locomotor state, even functioning at rest. Taken together with their extensive supraspinal input^38^, we suggest that the V3 neurons can be more generally viewed as spinal cord command neurons that regulate the gain of the motor system.

## Results

### V3 neurons widely innervate lumbar motoneurons and premotor neurons

Considering that silencing of V3 neurons leads to a marked loss in the ability of the isolated spinal cord to generate strong locomotor-like activity^30^, we first explored whether V3 neurons directly increase activity of motoneurons and premotor neurons, acting as a gain control mechanism. To this end, we started by examining the anatomy of V3 innervation of motoneurons throughout the lumbar spinal cord. To accomplish this, we generated *Sim1^Cre+/-^; Ai34^(RCL-Syp/tdT)-D^*mice expressing synaptophysin-Tdtomato specifically at V3 axon terminals, to visualize V3 presynaptic boutons (Figure 1A). A subsequent cross with *Hb9^GFP^* mice resulted in *Sim1^Cre+/-^; Ai34^(RCL-Syp/tdT)-D^; Hb9^GFP^* mice allowing for an examination of V3 synapses onto motoneurons (Figure 1A), where GFP labels motoneurons (Figure 1B)^41^. We found that V3 neurons established extensive synapses with both the soma and dendrites of motoneurons throughout the ventral horn (Figure 1B).

**Figure 1.**
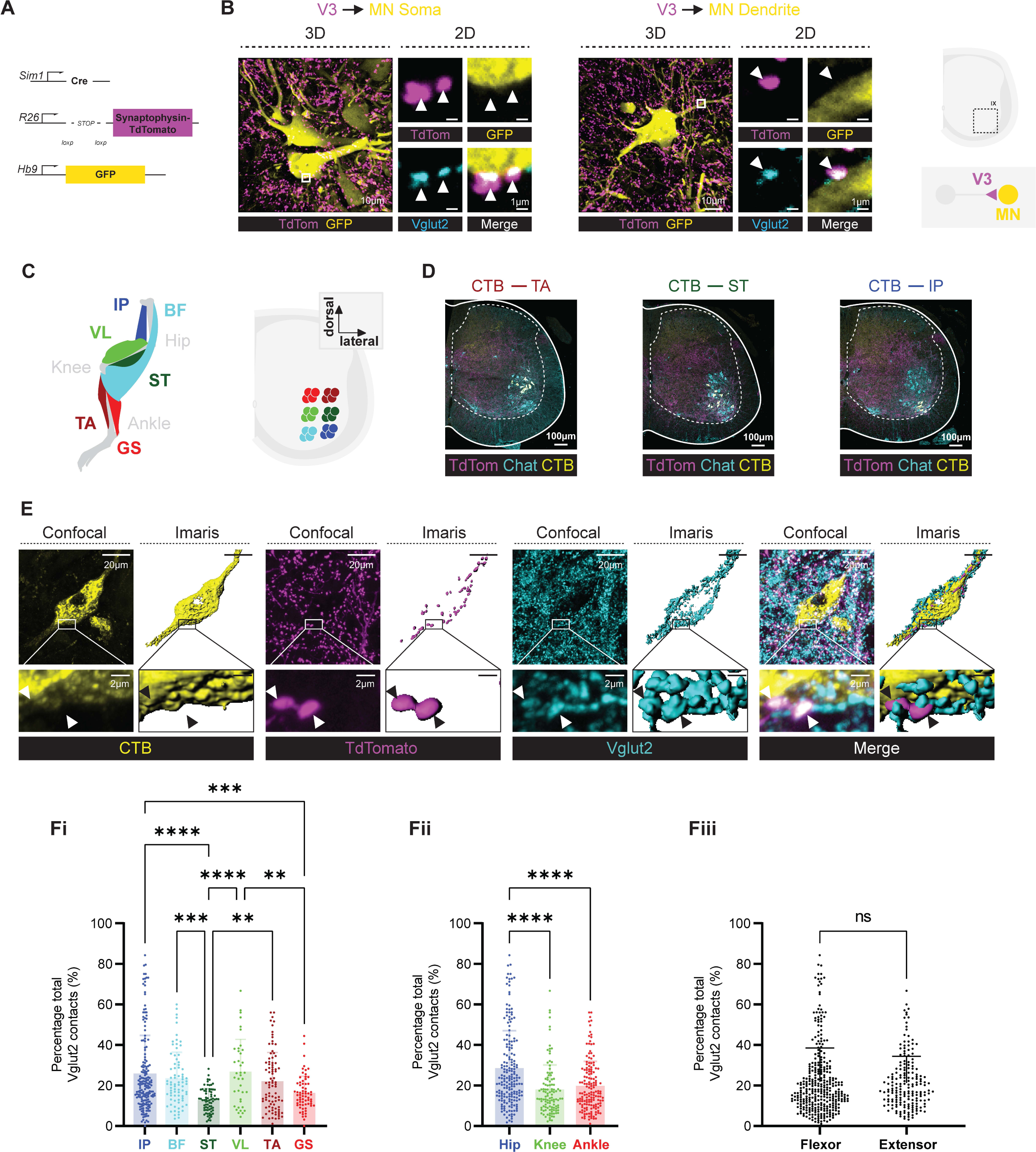
V3 neurons widely innervate lumbar motoneurons. A. Diagram of genetic strategy to generate *Sim1^Cre+/-^; Ai34^(RCL-Syp/tdT)-D^; Hb9^GFP^* mice. *Sim1^Cre+/-^* mice were crossed with *Ai34^(RCL-Syp/tdT)-D^* mice containing Synaptophysin TdTomato downstream of a loxP-flanked STOP cassette to generate *Sim1^Cre+/-^; Ai34^(RCL-Syp/tdT)-D^* mice. A subsequent cross with *Hb9^GFP^* mice resulted in *Sim1^Cre+/-^; Ai34^(RCL-Syp/tdT)-D^; Hb9^GFP^*mice. B. In *Sim1^Cre+/-^; Ai34^(RCL-Syp/tdT)-D^; Hb9^GFP^* mice, V3 axon terminals make glutamatergic contacts onto GFP expressing motoneuron somata (left) and dendrites (middle), as illustrated on the right. C. Cholera toxin subunit B (CTB) conjugated with Alexa FluorTM 488 was injected into various hindlimb muscles, as shown in the diagram (left), that retrogradely traced the motoneuron pools (right) innervating corresponding colour-coded muscles. D. Representative images of CTB positive motorneuron pools that innervate tibialis anterior (TA) (left), semitendinosus (ST) and iliopsoas (IP) muscle (yellow). Chat-immunolabelled motoneurons (Green) verify the CTB retrograde labelling. TdTom (red) labelled synaptic terminals of V3 neurons. E. IMARIS 3D reconstruction of retrogradely labelled CTB motoneuron, TdTomato labelled V3 pre-synapse terminal and immunolabelled Vglut2 labelled all excitatory synapses on motoneurons. F. V3 synaptic contacts on individual motoneurons. (i) Percentage of V3 contacts relative to the total VGLUT2^+^ contacts inputs on individual motoneurons in different motoneuron pools. N number of MNs counted for each muscle: iliopsoas = 178 cells, semitendinosus= 72 cells, tibial anterior= 88 cells, bicep femoris= 79 cells, vastus lateralis= 39 cells, medial gastrocnemius= 68 cells. N number of animals counted for each muscle: iliopsoas = 5 animals, semitendinosus= 3 animals, tibial anterior= 3 animals, bicep femoris= 3 animals, vastus lateralis= 1 animals, medial gastrocnemius= 2 animals. (ii - iii) same data as (i) but grouped by muscles innervating hip, knee, and ankle muscles (ii) or by function (iii). Significant differences indicated with **, 0.001 ≤ P < 0.01; ***, 0.0001 ≤ P < 0.001; ****, P < 0.0001, computed Tukey HSD test for multiple comparisons.

To delineate which motoneuron pools were innervated, we injected Cholera toxin subunit B (CTB) conjugated with Alexa Fluor 488 into various hindlimb muscles of *Sim1^Cre+/-^; Ai34^(RCL-Syp/tdT)-D^*mice (Figure 1C & D). Employing confocal microscopy and IMARIS 3D reconstruction, we observed that V3 neurons extensively innervate all 6 investigated motoneuron types that control muscles of the hip, knee, and ankle, which are widely distributed from rostral to caudal across the entire lumbar spinal cord. Remarkably, 15 – 25% of all the glutamatergic interneuron input (VGLUT2^+^) to motoneurons arose from V3 neurons (Figure 1E & F). Comparing flexor and extensor muscles, there was no difference in total motoneuron innervation (Figure 1Fiii). However, comparing across functional groups, the hip flexors-extensors received more V3 input than those of the knee or ankle (40.6% or 27.5% more, respectively, Figure 1Fii) (Supplementary Table). The hip and knee extensors, bicep femoris (BF) and vastus lateralis (VL), and the hip and ankle flexors, iliopsoas (IP) and tibialis anterior (TA) received the most V3 input (not significantly different between these muscles), whereas the knee flexor, semitendinosus (ST), and ankle extensor, gastrocnemius (GS), received less V3 input (Figure 1Fi).

In addition to innervating motoneurons, we observed that V3 neurons recurrently innervated other V3 neurons. To visualize this, we crossed *Sim1^Cre+/-^; Ai34^(RCL-Syp/tdT)-D^* mice with E*GFP^RCE:loxP^* resulting in *Sim1^Cre+/-^; Ai34^(RCL-Syp/tdT)-D^; EGFP^RCE:loxP^* mice, facilitating the observation of V3 cytoplasm in green and synapses in red (Figure 2Ai). We observed V3 synapses on both the cell body and dendrites of all other V3 neurons examined (Figure 2Aii), including both medial and lateral subpopulations of ventral V3 neurons (Figure 2C).

**Figure 2.**
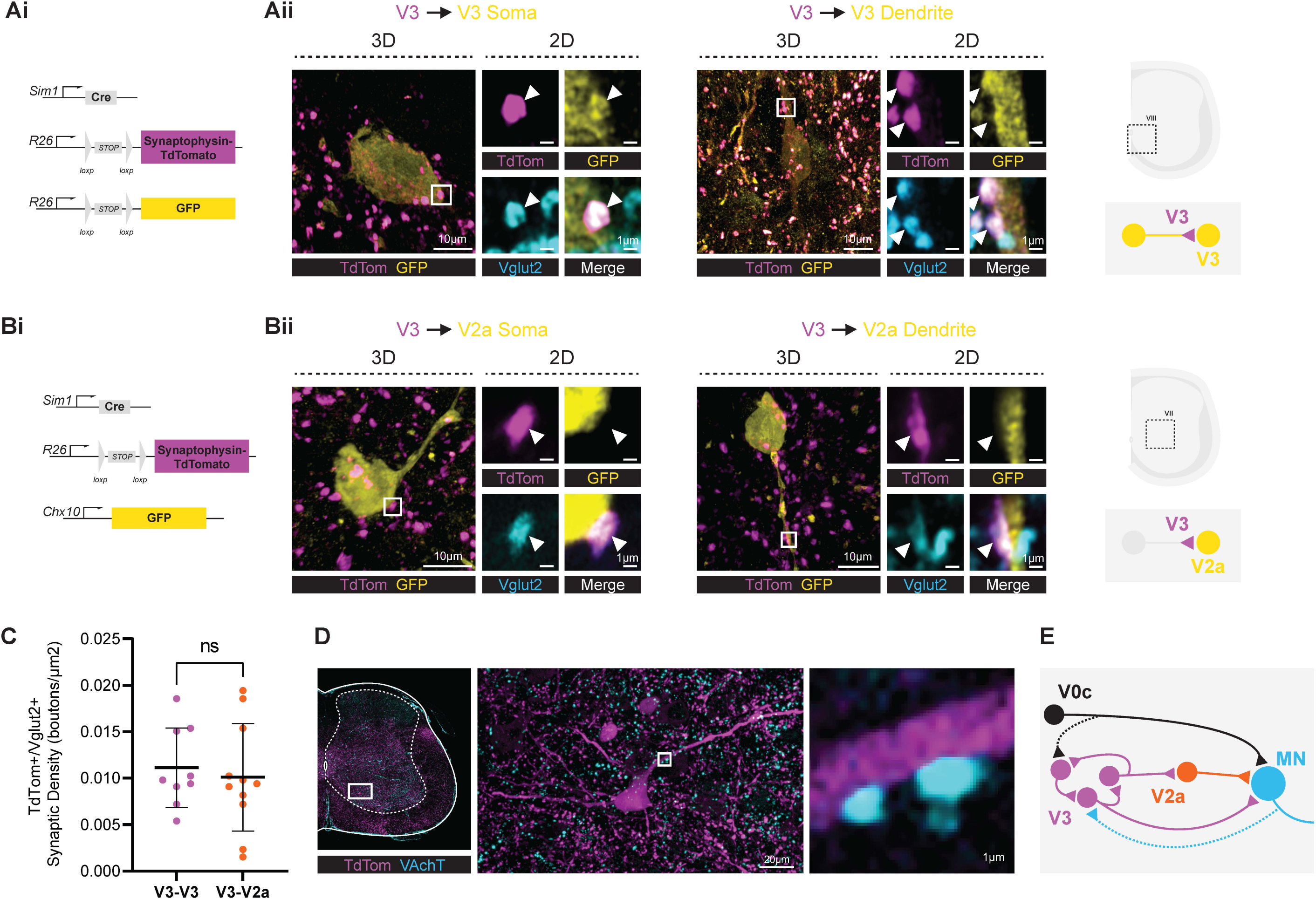
V3 neurons broadly innervate lumbar premotor neurons. A. V3 neurons make contacts with V3 neurons. (i) *Sim1^Cre+/-^* mice were crossed with *Ai34^(RCL-Syp/tdT)-D^* and *Tau^GFP^*mice containing Synaptophysin TdTomato and GFP downstream of a loxP-flanked STOP cassette to generate *Sim1^Cre+/-^; Ai34^(RCL-Syp/tdT)-D^; Tau^GFP^* mice that enabled the observation of V3 cytoplasm in green and synapses in red. (ii) Representative images showing V3 excitatory projections (V3-synaptophysin, magenta; Vglut2, blue) to the soma and dendrite (GFP, yellow) of other V3 neurons, which was similar in n = 5 V3 neurons from throughout the ventral horn of 5 mice. (iii) Schematic diagram shows the region where images were taken from. B. V3 neurons form connection to V2a neurons. (i) *Sim1^Cre+/-^; Ai34^(RCL-Syp/tdT)-D^* mice were crossed with *Chx10 ^EGFP^* mice to generate *Sim1^Cre+/-^; Ai34^(RCL-Syp/tdT)-D^;Chx10 ^EGFP^* mice to examine the V3-V2a connection. (ii) Representative images showing that V3 neurons form excitatory projections (synaptophysin, red; Vglut2, green) to the soma and dendrite (yellow) of V2a neurons, which was similar in n = 5 V2a neurons from throughout the ventral horn of 5 mice (iii) Schematic diagram shows the region where images were taken from. C. Percentage of V3 contacts relative to the total VGLUT2^+^ contacts inputs on V3 neurons and V2a neurons. N number of V3 cells counted = 9, N number of V2a cells counted = 1. From 1 animal of each. D. V3 neurons receive cholinergic innervations. In *Sim1^Cre+/-^; Rosa^.lsl.tdTom^*mice, TdTomato labelled V3 neurons. Immunolabelled VAchT to show cholinergic innervations. Left image shows the region where the image was taken from. Middle and right images show the enlarged areas of interest. E. Schematic diagram shows the summary of the premotor neuron circuit that V3 neurons form.

Interestingly, V3 neurons also projected to other excitatory interneurons, further increasing their action. That is, projections of V3 neurons to V2a neurons (Chx10^+^) were observed, as evidenced by V3 synaptophysin contacts on EGFP^+^ cells in *Sim1^Cre+/-^; Ai34^(RCL-Syp/tdT)-D^; Chx10^eGFP^*mice, while reciprocal synapses from V2a neurons to V3 neurons were not detected (Figure 2B). In addition, using *Sim1^Cre+/-^; Ros^.lsl.tdTom^* mice, we confirmed that V3 neurons also receive cholinergic innervations, which could be from either motoneurons or V0c neurons or both, to generate recurrent excitation (Figure 2D). Overall, these results demonstrate that V3 neurons form an extensive feedforward excitatory circuit, both exciting motoneurons and premotor neurons (Figure 2E).

### Activation of lumbar V3 neurons broadly enhances motoneuron and muscle activity

To explore whether the widespread direct and indirect innervation of motoneurons by V3 neurons could rapidly increase motor output, we next employed ChR2 to optogenetically activate V3 neurons in *Sim1^Cre+/-^; Ai32^RCL-ChR2(H134R)^*^EYFP^ mice (abbreviated V3ChR2 mice). Isolating neonatal P2-3 spinal cords from V3ChR2 pups, we positioned them ventrally upward in a perfusion chamber, and applied suction electrodes for electroneurogram (ENG) recordings on the lumbar ventral roots innervating extensor and flexor muscles (Figure 3Ai & Bi). We observed that localized light activation of V3 neurons at just one segment (either the L2 or L5) elicited robust motor responses across all lumbar segments, from L2 to L5 (Figure 3Aii, iii & Bii, iii), which likely included varying degrees of extensor and flexor motoneuron responses, because the L5 root is extensor dominated and L2 flexor dominated^42^. Unexpectedly, the activation of just the right V3 neurons at L2 or L5 rapidly evoked motor activity bilaterally (Figure 3Aii, iii & Bii, iii), and thus we examined the origin of this by grading the intensity of light applied at L2. At the lowest light intensity near threshold (T, as used in Danner et al., 2019^39^), the contralateral ventral roots exhibited a very short latency response consistent with a monosynaptic pathway of commissural motoneurons innervating contralateral motoneurons^37^, whereas the ipsilateral roots at the same segment had longer polysynaptic latency responses (∼2 ms later). However, increasing light intensity activated both contralateral and ipsilateral motoneurons at a similar short time, suggesting that deeper more dorsal V3 neurons or higher threshold axon tracts with ipsilateral actions were activated by light (see Discussion).

**Figure 3.**
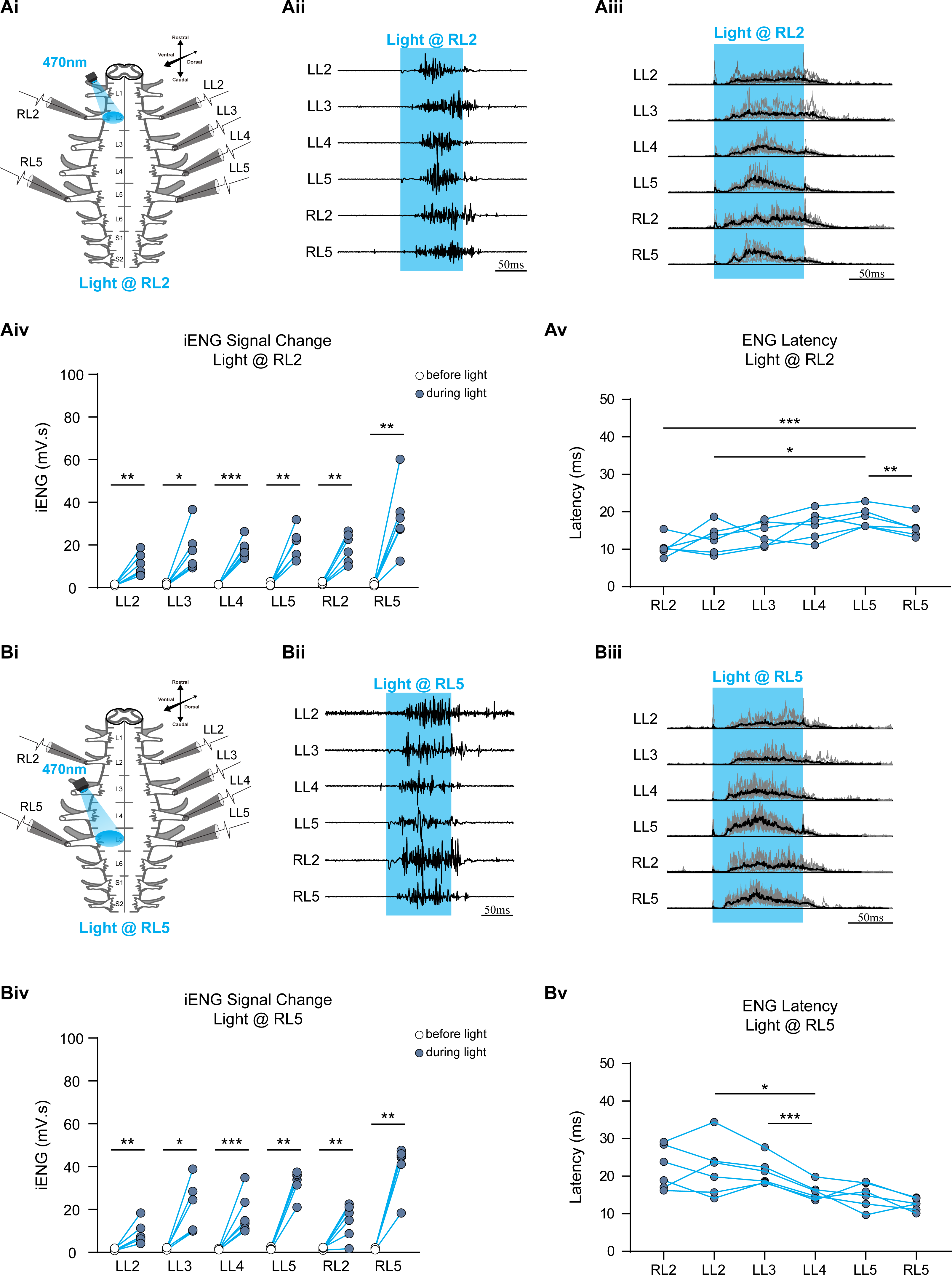
Activation of local V3 neurons broadly enhances motor output in the isolated neonatal lumbar spinal cord. A. Optical activation of L2 V3 neurons evokes activities in all lumbar ventral nerve roots. (i) Experimental setup. Isolated spinal cord from neonatal P2-3 V3ChR2 pups was positioned ventral side up in a recording chamber, and suction electrodes for electroneurogram (ENG) recordings were applied on the lumbar ventral roots. Focal light activation of V3 neurons at right L2 region. (ii-iii) Optical activation of right L2 (at 4xT, T, threshold for light response) elicited robust motor responses across all lumbar segments, from L2 to L5 on the left side and L2 and L5 on the right side. The blue shaded area represents the optical stimulation period. Representative raw (ii) and averaged rectified (iii) signals are shown. Black thick lines are the averaged traces of multiple averaged rectified ENG recordings from each animal (n = 6 animals, each animal has 10-15 trials, light gray lines in iii). (iv) The mean integral ENG during drug-induced locomotor-like activity (n = 6 mice) before (white circles) and during (blue circles) optical activation on L2. Each circle represents the average value of the estimated parameter for one animal. Light blue lines connecting two circles represent the parameter increase in one cord. Significant differences indicated with ∗, P < 0.05; ∗∗, 0.01 < P < 0.05; ∗∗∗, 0.0001 < P < 0.001, paired t-test. (v) The latency delay between light onset and ENG activity was compared. Significant differences indicated with ∗, P < 0.05; ∗∗, 0.01 < P < 0.05; ∗∗∗, 0.0001 < P < 0.001, paired t-test. Liner regression for the order of RL2, LL2-5, RL5, slope= 0.001292, R^2^= 0.3273, slope is significantly non-zero, P= 0.0003. B. Optical activation of V3 neurons at right L5 region. Similarly to the described experimental setup in A (i) with ENG recordings at lumbar ventral roots of isolated neonatal spinal cords, but the focal light activation of V3 neurons is at right L5 region. (ii-iii) Optical activation of right L5 elicited robust motor responses across all lumbar segments (n = 6 animals, each animal has 10-15 trials, light gray lines in iii). (iv) The mean integral ENG during drug-induced locomotor-like activity (n = 6 mice) before (white circles) and during (blue circles) optical activation on L5. Each circle represents the average value of estimated parameter for one animal. Light blue lines connecting two circles represent parameter increase in one cord. Significant differences indicated with ∗, P < 0.05; ∗∗, 0.01 < P < 0.05; ∗∗∗, 0.0001 < P < 0.001, paired t-test. (v) The latency delay between light onset and ENG activity was compared. Significant differences indicated with ∗, P < 0.05; ∗∗∗, 0.0001 < P < 0.001, paired t-test. Liner regression for the order of RL2, LL2-5, RL5, slope= -0.002144, R^2^= 0.4313, slope is significantly non-zero, P<0.0001.

To quantify this enhanced motoneuron activity with optogenetic V3 activity, we measured the integrated ventral root ENG (iENG) 20 ms before light and after first ENG activity onset during light exposure (Figure 3Aiv & Biv). The results indicate a significant rapid elevation in motoneuron activity for each nerve, suggesting that V3 neuron activation in either segment could induce rapid extensive global bilateral excitation within the lumbar motoneurons. Notably, when L2 lumbar V3 neurons were stimulated, the ENG activity latency of the L5 nerve was significantly longer than that of the L2 nerve (Figure 3Av), and more generally the latency increasing proportional to the distance from the stimulation site to the recorded ventral root (Figure 3Av). Overall, these findings suggest that V3 neurons within the L2 or L5 segment are capable of rapidly initiating bilateral motoneuron excitation throughout the lumbar motor circuit.

We next investigated in the awake adult V3ChR2 mice whether localized activation of V3 neurons also evoked widespread motoneuron excitation. By applying light through a window implanted over the L2 spinal cord (Figure 4A, bilateral light), we observed that activation of V3 neurons in one lumbar spinal segment elicited muscle activity across all joints of the hindlimb, and thus activating motoneurons throughout the lumbar spinal cord (Figure 4B). Interestingly, despite the anatomical connection of V3 neurons to all motoneurons, the light evoked electromyogram (EMG) in extensor muscles was larger than in flexors (Figure 4B). Even when we normalized each EMG to its peak observed during voluntary activity (e.g. walking), to account for inter-animal variability, the extensor EMG remained larger than flexor EMG (Figure 4Ci & ii). Furthermore, the latency of EMG from light onset was very short in extensor muscles, likely reflecting the monosynaptic connections to motoneurons from V3 neurons, whereas the flexor latency was slower in onset (Figure 4Ciii). The observed fast widespread activation of motoneurons only occurred when the light was sufficient to penetrate deep into the spinal cord, 10x the light threshold to produce long latency more localized responses, and thus likely involved the ventral V3 neurons (see Discussion).

**Figure 4.**
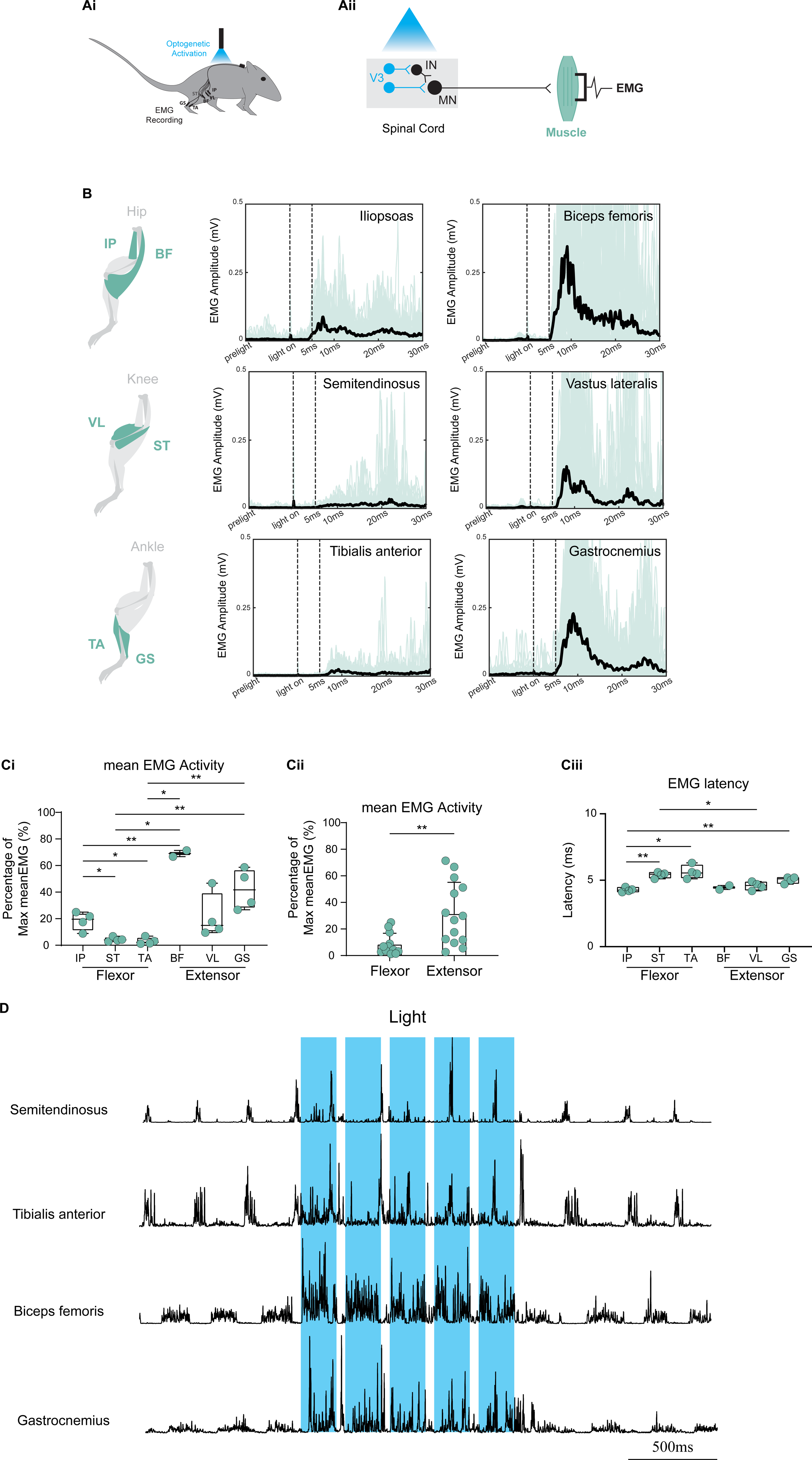
Activation of lumbar V3 neurons broadly enhances muscle activity. A. The schematic diagram illustrates the experiment protocol. (i) Experimental setup. (ii) A window was implanted over the L2 spinal cord (or sometimes L4) to apply light activating V3 neurons in V3ChR2 adult mice. Muscle activities are recorded by the implantation of fine electrodes. B. Optical activation of V3 neurons in one lumbar spinal segment with a brief light pulse (15 ms, 450 nm, 10.4 mW/mm^2^ pulse; at 10xT, where T is the response threshold) elicited muscle activity across all joints of the hindlimb, including hip (IP and BF), knee (ST and VL) and ankle (TA and GS). Black thick lines are the averaged traces of all original rectified EMG recordings (light green lines) from different mice (n =4 for IP, ST, VL, TA and GS, and n=2 for BF). C. Activation of V3 neurons in one lumbar spinal segment elicited rapid and extensor biased muscle activity. (i) Plot of the mean EMG response amplitude relative to maximal voluntary EMG for animals of B. Same data as (ii), but muscles grouped into flexor and extensors. (iii) Plots of latency of rectified EMG response to light and mean EMG response amplitude relative to maximal voluntary EMG, for animals of B. N number of animals counted for each muscle: iliopsoas = 4 animals, semitendinosus= 4 animals, tibial anterior= 4 animals, bicep femoris= 2 animals, vastus lateralis= 4 animals, medial gastrocnemius= 4 animals. Significant differences indicated with ∗, P < 0.05; ∗∗, 0.01 < P < 0.05, paired t-test. D. Rectified raw EMG recordings show that focal optogenetic activation of V3 neurons also increases muscle activities during walking on the treadmill. Blue shaded area is the optical stimulation period. Similar responses in n = 3/3 mice.

This widespread extensor-biased activation of muscles by focal optogenetic activation of V3 neurons occurred both when the mouse was resting (Figure 4) or walking (Figure 4D), indicating that the V3 circuit that generates extension does not depend on the state of the CPG. Overall, these results suggest that V3 neurons rapidly excite all motoneurons, but there is an extensor muscle bias that might help produce weight support during walking.

### V3 neurons cause sustained motoneuron activity

Upon administering a single short 10-15 ms light pulse to V3 neurons in awake V3ChR2 mice, there was a rapid sustained increase in muscle activity that persisted 50 – 100 ms beyond the light exposure period (Figure 5Ai). A longer, 150 ms light exposure elicited an even longer sustained increase in muscle activity (Figure 5B). This phenomenon was consistently observed across different muscle groups, including the IP, ST, TA, BF, VL and GS. To further quantify this sustained muscle activity, we measured EPSPs evoked in motoneurons by brief light pulses in vitro in the adult spinal cord from V3ChR2 mice (gold in Figure 5A) and compared them to the EMG from awake mice. Overall, the duration of the EMG evoked by a light pulse was similar to the EPSP duration, though we noted that the EMG was composed of bursts. To examine whether these bursts trivially resulted from synchronization of motoneuron firing by the light, we picked single motor units out of the gross EMG records (5Aiii). As expected, the firing of the single motoneuron spikes after the light onset was at the burst interval of about 10 – 20 ms, suggesting that the motoneurons were simply responding to the long EPSP with 50 -100 Hz repetitive firing that was synchronized to the EPSP onset (see method of Norton et al., 2008)^43^. Overall, these results suggest that not only do V3 neurons have fast direct connections, but they are also intrinsically capable of producing long-lasting activity to sustain motor output, as shown by Lin et al. 2022^44^.

**Figure 5.**
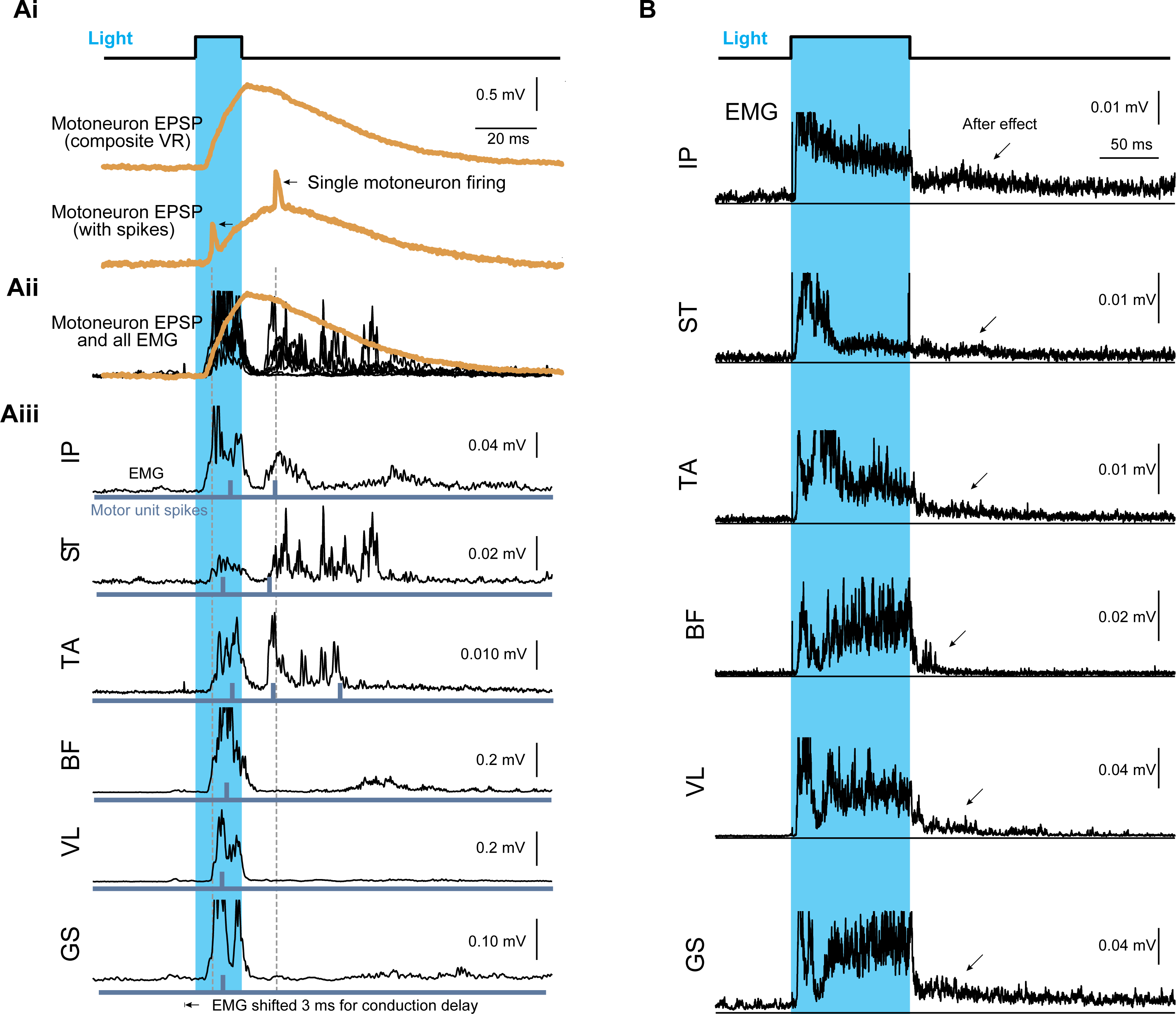
V3 neurons cause sustained motoneuron activity. A. (i) Administering a single short light pulse to V3 neurons in vitro in the adult spinal cord of ChR2 mice (gold lines) rapidly evoked a 50 – 100 ms long composite EPSP recorded from motoneurons of the ventral roots with a grease gap method, with sometimes individual motoneuron spikes visible (lower trace). (ii) The same short light pulse in awake V3ChR2 mice evoked a similar sustained increase in all muscle activity (black), with overlayed EPSP from (i) shown. (iii) Individual muscle responses. A single motor unit is picked from the gross EMG records, as shown by the gold spike in (i) and the blue bar in (iii). The blue shaded area represents the optical stimulation period. Similar responses in all mice from Fig 4, and n = 5/5 mice recorded in vitro. B. 150ms light exposure elicited an even longer sustained increase in all muscle activities. The blue shaded area represents the optical stimulation period. The after-effects are indicated by arrows. Similar responses in n = 5/5 mice.

### V3 neurons are necessary for increasing muscle activity with locomotor intensity

Finally, to examine whether V3 neurons are necessary for controlling the gain of motoneuron output, we genetically silenced them by crossing *Sim1^cre+/-^* mice with *VGlut2^flx/flx^* mice to stop glutamate transmission (V3OFF mice). These mice, without functional V3 neurons, were able to function relatively normally, but they were slow^35^, clumsy (with wide meandering stepping; as in Zhang et al., 2008)^30^, and only explored a new cage for an unusually short period (n = 10/10 mice). This seemed to be due to an overall weakness in these mice, but the weakness proved hard to quantify because V3OFF mice have intact voluntary supraspinal control and accordingly adopt compensatory behaviours, like rarely fully relaxing, so muscles are partly pre-activated in case they have to move. Thus, to quantify whether V3OFF mice were weaker, we challenged them with locomotor tasks of increasing intensity.

When we forced V3OFF mice to walk on a level treadmill with increasing speed, they failed reach the high speeds normal wildtype mice can reach (∼<20 m/s for V3OFF mice with EMG implants, vs <80 cm/s in wildtype mice; Zhang et al., 2019)^35^, and when the treadmill was inclined 17.5°, they failed to keep up with the slow treadmill speed and so gradually drifted to the bottom of the treadmill, essentially walking slower than wildtype mice. Furthermore, when placed in water, they swam with a slower forward velocity with rather weak propulsive motions, compared to in wildtype mice (see Supplementary Video). Considering the difficulties in comparing absolute EMG amplitude between animals, we normalized activity in the more intense locomotion by that in slow level ground walking to further compare the activity in wildtype and V3OFF mice. In wildtype mice, most muscle EMGs increased markedly during these more demanding tasks, especially in the extensor muscles (∼doubling) (Figure 6B & C). However, in V3OFF mice, these increases were much less or absent, significantly lower in extensor muscles and the hip flexor muscle, IP. Overall, the muscle activity with functional V3 neurons was dramatically higher than without, by ∼ 50 – 400%, depending on the muscle (Figure 6C), consistent with a generalized weakness in V3OFF mice.

**Figure 6.**
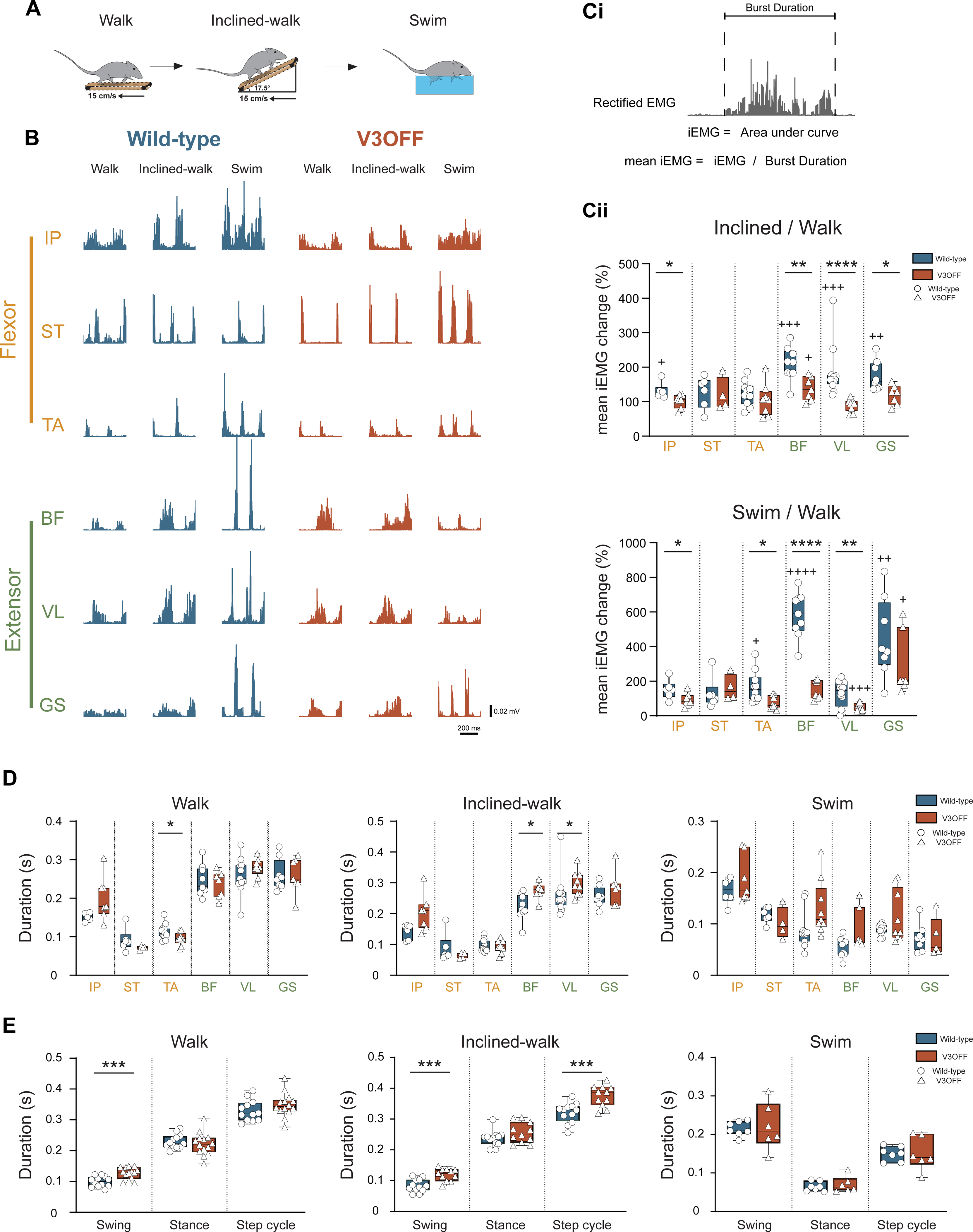
V3 neurons are necessary for increasing muscle activity with locomotor intensity. A. The schematic illustration of the experimental setup. B. Representative rectified raw EMG traces of WT (left) and V3OFF (right) mice during level-walking, inclined-walking, and swimming, with two complete locomotor cycles shown for each. C. V3OFF mice had difficulties increasing muscle activities in intense locomotion. (i) The method and equation to measure the mean of the iEMG from one step. (ii) Box mice of means of the iEMG increase of each muscle in WT and V3OFF mice between level-walking (walk) and inclined-walking (inclined-walk) (top), and between level-walking (walk) and swimming (swim) (bottom). The blue and red boxes indicate the 5 to 95 percentile of data distribution. The means of each mouse are shown as white circle for wildtype and as white triangle for V3OFF. Significant difference in V3OFF compared to wildtype indicated with * and significant change in EMG compared to level ground walking indicated with +, computed with an unpaired t-test, 0.01 ≤ P < 0.05; **, 0.001 ≤ P < 0.01; ****, P < 0.0001, for same mice as in C. D & E. V3OFF mice have little change in muscle activity burst duration and step phase duration. Box plots, means of the activity burst duration of each muscle (D) and step phase duration (E) in WT and V3OFF mice between level-walking (walk, left), inclined-walking (inclined-walk, middle), and swimming (swim, right). The blue and red boxes indicate the 5 to 95 percentile of data distribution. The means of each mouse are shown as white circle for wildtype and as white triangle for V3OFF. Significant differences indicated with *, 0.01 ≤ P < 0.05; ***, 0.0001 ≤ P < 0.001, t-test. 13 wildtype and 11 V3OFF animals. N number of muscles in wildtype: IP=6, ST=6, TA=10, BF=9, VL=12, and GS=8; in V3OFF: IP=7, ST=4, TA=8, BF=7, VL=9, and GS=7.

In contrast to their marked weakness, V3OFF mice had comparatively less difficulty in regulating the timing of EMG during walking and swimming. In each of these three tasks, burst durations in most muscles of V3OFF mice were not different or changed < 30% compared to wildtype mice (Fig 6D). Specifically, of all the hindlimb muscles only the hip (IP) and ankle (TA) flexors, increased slightly in burst duration during level walking in V3OFF mice, only the hip (BF) and knee (VL) extensors increased in burst duration during inclined walking, and no muscle changed in duration during swimming (Figure 6D). Furthermore, examining the timing of the locomotion, we found small changes in V3OFF mice. That is, in V3OFF mice the locomotor cycle duration was unchanged in slow level walking or swimming and only increased ∼20% in inclined walking, the swing (protraction) phase duration was only increased ∼20% in two walking behaviours and unchanged in swimming, and the stance (retraction) phase was unchanged in all forms of walking and swimming (Figure 6E). Overall, with a loss the functional V3 neurons, the relative lack of changes in timing, compared to the marked changes in EMG amplitude without functional V3 neurons points to a crucial role of V3 neurons in amplifying the motoneuron output, as well as an ability for mice to compensate for the loss of these important neurons (see Discussion).

## Discussion

Our results demonstrate that V3 neurons widely innervate hindlimb motoneurons and premotor neurons throughout the lumbar spinal cord, providing a remarkable 15 - 25% of the total interneuron input to motoneurons (VGLUT2^+^). Together with their long propriospinal tracts, this allows the activation of V3 neurons at just one segment to rapidly produce sustained motoneuron activity across the entire lumbar spinal cord. Interestingly, despite similar number of V3 inputs onto extensor and flexor motoneurons, activation of V3 neurons functionally produces much more activity in extensor motoneurons, providing extensor tone in both the walking or sitting mouse. Finally, we show that V3 neurons are necessary for augmenting force production during locomotion, without which mice are weak and slow, unable to augment extensor force during demanding locomotor tasks, like climbing or swimming (in V3OFF mice), though they still remain capable of regulating the timing of locomotion. Overall, our results show that V3 neurons augment motoneuron output across the repertoire of mouse behaviours, providing a direct pathway for generating muscle force that regulates motoneuron output gain both during resting and locomotor states.

### Extensor motoneuron bias with V3 neuron activation

A key issue that remains unresolved is why there is an extensor bias in the V3 neuron responses, despite the anatomy suggesting otherwise (also see Danner et al., 2019^39^). One possibility is that there are functional subpopulations of V3 neurons and their premotor neuron targets that specifically innervate either extensor or flexor motoneurons, and perhaps extensor-related V3 neurons are more dominant under the conditions we studied. Interestingly, there does not appear to be a task-dependence or motoneuron-activity-dependence to the extensor bias because we see it when animals are quietly sitting or when they are walking, ruling out the simple explanation that a lack of excitability of the flexor motoneurons leaves them too far below threshold to respond forcefully. This makes it even more puzzling why V3 activation only weakly activates flexor motoneurons, suggesting that there is something that actively inhibits flexor-related V3 neurons. Possibly, the recently described recurrent inhibition of V3 neurons might allow extensor V3 neurons to recurrently inhibit flexor V3 neurons to assure reciprocal muscle control^44^.

### Rapid and sustained motoneuron activation by V3 neurons

Another issue is explaining why the muscle responses to brief V3 activation are so fast and long lasting. Certainly, the monosynaptic connections from V3 neurons to motoneurons ensure rapid activation, but the more slowly rising, larger and longer lasting overall response is unlikely to be due to this monosynaptic connection alone. Instead, the recently described persistent sodium currents in V3 neurons may provide an amplification and prolongation of the V3 neuron activity^44^. Furthermore, our finding that V3 neurons recurrently excite themselves and other premotor neurons (V2a neurons; Figure 1), as well as receive recurrent excitation from motoneurons^36^, suggests that this microcircuit amplifies and prolongs V3 neuron function. Together, the sustained input to motoneurons from V3 intrinsic properties and microcircuits is likely to engage persistent sodium and calcium currents in motoneurons themselves, furthering amplify their output^45–48^.

### Bilateral motoneuron activation by V3 neurons

Considering that commissural axons account for almost all the connections from V3 neurons to motoneurons (∼85%, likely ventral V3 neurons, some Olig3+^30,34^), our observation that moderate light activation of V3 neurons on just one side of the L2 lumbar cord rapidly activates widespread motoneurons on both sides of the whole lumbar cord is puzzling (also see Danner et al.^39^). Perhaps this strong bilateral activation suggests that V3 neurons rapidly excite motoneurons by activating premotor neurons as well as motoneurons, including by the connections to other V3 or V2a neurons that we have demonstrated. Because the vast majority of V3 neurons only send axons across the midline, V3 activity on one side of the cord should quickly lead to V3 activity on the other, with ultimately a large bilateral motoneuron activation (similar to the model in Danner et al., but directly to V3 neuron). Moreover, ipsilaterally projecting V3 neurons may also contribute to motoneuron input, but these only account for about 5% of V3 neurons by postnatal stages (Onecut2+)^34^, and it is observed that while local ipsilateral projections occur from the most medial ventral V3 neurons to nearby V3 neurons, the remaining V3 neurons only have inputs from elsewhere, most likely from across the midline. Functionally, V3-to-V3 activation may well ricochet down the spinal cord and include both crossed and uncrossed connections, accounting for some of the activity delay we see with distance down the cord, which is especially evident in the room temperature neonatal spinal cord preparation where synaptic and conduction delays are long(∼ 2.5 – 3 ms synaptic latency) ^49,50^.

An additional explanation for why local light application in V3ChR2 mice evokes rapid bilateral motoneuron output arises from considering the mechanism by which ChR2 excites V3 neurons. We have previously shown that the depolarization produced by ChR2 is generally not sufficient alone to produce postsynaptic potentials on the V3 neuron targets, but relies on the sodium channel, since this action is blocked by TTX^44^. This implies that for light activation of V3 neurons to evoke responses in their target motoneurons, sufficient ChR2 must be located near a spike initiation zone and ultimately initiate a spike that is transmitted to the terminals. Considering that ChR2-EYFP covers most of the V3 neuron membrane (data is not shown), there is plenty of ChR2 available in the soma and proximal dendrites to activate the sodium channels in the axon initial segment (as indicated by Figure1A in Danner et al., 2019)^39^. Hence, light application on one side of the cord is most likely to initiate a spike at the V3 soma on that side, as we have implicitly assumed in our discussion above. However, we cannot rule out light initiating a spike by ChR2 located on axons arising from contralateral V3 neurons. Thus, light applied to one side of the cord may produce bilateral contralateral and ipsilateral motoneuron activation via ChR2 on the soma of ipsilateral V3 neurons and crossed axon from contralateral V3 neurons, respectively. Settling this issue will require future studies with focal light activation of individual soma or axons.

### Connectivity and dynamic activation patterns of V3 neurons

The question of which subpopulation of V3 neurons innervates motoneurons also remains unclear. Considering that mouse ventral V3 neurons closely resemble cat lamina VIII commissural neurons of Jankowska^37,38^, it is reasonable to suppose that these two populations may be the same, allowing us to incorporate findings from these earlier cat studies. That is, like ventral V3 neurons, these cat neurons are in lamina VIII, commissural, and widely innervate contralateral motoneurons. Furthermore, Jankowska found that these laminae VIII neurons are the only commissural neurons that innervate motoneurons and virus-tracing data from Ronzano et al. showed commissural premotor neurons mainly locate in laminae VIII^51^. Laflamme et al. also showed that exclusive commissural V3 neurons are directly involved in the amplitude regulation of crossed reflex^52^. Therefore, it is likely that mouse V3 neurons in laminae VIII (ventral V3) are the major subpopulation that innervates motoneurons. Consequently, our finding that light applied at L2 rapidly activates motoneurons along the whole lumbar spinal cord (from IP at L1-L2 to GS at L5-L6) suggests that ventral V3 neurons form long descending and ascending commissural axon tracts with branches that innervated motoneurons along the whole lumbar cord (these are likely ventrally located tracts). This is consistent with our observation that while light applied dorsally activates motoneurons rapidly, it did so at high light intensities many time that to produce slower responses, and thus likely activates ventral V3 neurons (see next paragraph). However, the detailed anatomy of these neurons awaits further study. Finally, it is also interesting that the cat laminae VIII neurons receive extensive sensory and supraspinal input, consistent with our limited understanding of mouse V3 inputs^38,52^. The input from vestibulospinal and reticulospinal tracts is particularly telling, as these are crucial for posture and locomotion.

If we take for granted the conclusion that only ventral V3 neurons innervate motoneurons, then our very fast activation of motoneurons by light likely arises from the direct activation of ventral V3 neurons. However, how this occurs with the light applied in vivo to the dorsal surface of the thick lumbar spinal cord requires explanation. Firstly, it should be remarked that V3 neurons have extensive dendritic arbours that extend throughout the intermediate and deep dorsal spinal cord, and thus, light applied dorsally may only have to activate ChR2 on the most dorsal dendrites to activate the ventral V3 neurons. Secondly, we found that fast powerful activation of motoneurons required about ten times the light intensity needed to produce slow weak responses, suggesting that only a small fraction of the applied light actually reaches ventral V3 neurons, consistent with the expected sharp light attenuation with distance. Lastly, it is possible that the V3-to-V3 connections might allow fast activation of motoneurons. These connections highlight the intricacy of V3 neuron interactions and the potential for complex circuit dynamics that facilitate rapid and extensive motoneuron activation.

### V3 neurons function as command neurons that globally recruit motor behaviours

In addition to amplifying the gain of motoneurons, V3 neurons may also amplify the activity of the many other neurons they innervate, including V2a (Figure 2), V1^30^, GAD2+^44^, sensory^53^ and autonomic neurons^54^. This broader role in gain control is evident in our previous observations that sufficient optogenetic V3 neuron activation in the isolated neonatal spinal cord can initiate locomotor-like activity in the extensor phase and can reset the timing of the CPG^39^. The extent to which such strong V3 interactions with the CPG occurs in the adult mouse is unknown, though the subtle changes in adult locomotor timing that we see with V3 neuron silencing, suggests that they are not much needed for regulating the basic timing of locomotion, as also concluded for zebrafish swimming^55^. However, V3 neurons receive extensive sensory input, including group I proprioceptive input^44^, perhaps allowing them to relay sensory information to the CPG, which indirectly affects timing. This mechanism may explain why ankle and hip proprioceptors can so powerfully modulate the step cycle during locomotion^56–60^.

While our anatomical studies and optogenetic activation of V3 neurons provide clear evidence that these neurons are sufficient to augment motor output, their function in normal behaviour remains less certain. Using a permanent knockdown of V3 neuron function, we demonstrate here that without V3 neurons, mice are weaker, severely lacking the ability to augment muscle force when challenged with inclined ramps and swimming tasks. Because these mice make numerous voluntary compensations that allow them to survive and function normally over their life, the role of V3 neurons in less forceful behaviours was hard to quantify, and thus future studies with rapid optogenetically silencing of V3 neurons may further clarify this issue. Nevertheless, without V3 neurons mice have a clear phenotype associated with this weakness, including being slow, clumsy and sedentary. More generally, our results from both genetically silencing and activating V3 neurons indicate the importance of V3 neurons not just in activating motoneurons but also in amplifying motoneuron output from the CPG.

In summary, V3 neurons directly depolarize motoneurons and premotor neurons allowing them to rapidly generate muscle force, without which mice are weak and slow. By doing so, this brings motoneurons closer to threshold, allowing other CPG circuit elements to excite them more easily, and thus indirectly amplifying CPG or any motor output.

Although we have shown that ventral V3 neurons circuits are recruited in locomotion like walking and swimming (with CFOS)^33^, these circuits also bypass the CPG for locomotion since they are functional when mice are sitting^34^, and they are driven by supraspinal input. While it is a semantic issue of whether to solely include the V3 neurons as part of the CPG for locomotion, it might be prudent instead to view these V3 neurons as more like spinal cord command neurons, resembling CST or RST neurons, since they directly activate many spinal neurons, including motoneurons^30^, CPG neurons (V2a and V1)^30^, sensory neurons^53^ and autonomic neurons^54^, ultimately increasing the gain of motor circuit outputs.

## Methods and Materials

### Mice

Adult and neonatal mice expressing Sim1^Cre^ were used to study V3 neurons. The generation and genotyping of Sim1^Cre+/-^ mice were described previously by Zhang et al. (2008). To visualize the synaptic terminals of V3 neurons, we crossed *Ai34^(RCL-Syp/tdT)-D^* mice (Jackson Laboratory, Stock No. 012569) with Sim1^Cre+/-^ mice to generate Sim1^Cre+/-^; *Ai34^(RCL-Syp/tdT)-D^* mice. A subsequent cross with *Hb9^GFP^*(Jackson Laboratory, Stock No. 005029) mice resulted in *Sim1^Cre+/-^; Ai34^(RCL-Syp/tdT)-D^; Hb9^GFP^* mice for examination the V3 synapses onto motoneurons. To visualize of both V3 soma and synaptic terminals simultaneously, we crossed *Sim1^Cre+/-^; Ai34^(RCL-Syp/tdT)-D^*mice with *EGFP^RCE:loxP^* (Jackson Laboratory, Stock No. 032037-JAX) to generate *Sim1^Cre+/-^; Ai34^(RCL-Syp/tdT)-D^; EGFP^RCE:loxP^* mice. To visualize V3 neurons *Ai14^(RCL-tdT)-D^* mice (Jackson Laboratory, Stock No. 007908) were crossed with *Sim1^Cre+/-^* to generate *Sim1^Cre+/-^; Ai14^(RCL-tdT)-D^* mice (Blacklaw et al., 2015). For checking the projections of V3 neurons to Chx10-V2a neurons, we crossed *Sim1^Cre+/-^; Ai14^(RCL-tdT)-D^* with Chx10^eGFP^ (Mutant Mouse Resource and Research Centers, MMRRC No. 011391-UCD) (Chopek et al., 2021). *Sim1^Cre+/-^; Ai32^RCL-ChR2(H134R)^*^EYFP^ mice were generated by crossing *Sim1^Cre+/-^* with *Ai32^RCL-ChR2(H134R)^*^EYFP^ (Jackson Laboratory, Stock No. 024109). Conditional knock-out of VGluT2 in Sim1-expressing V3 neurons, Sim1^Cre+/-^; VGluT2 ^flox/flox^ (V3OFF mice) was described previously by Chopek et al. (2018). In all our experiments, we used male and female mice equally. All procedures were performed in accordance with the Canadian Council on Animal Care and approved by the University Committees on Laboratory Animals at Dalhousie University and University of Alberta.

### Spinal cord tissue dissection, processing, and sectioning for histology

Spinal cords were obtained at postnatal stages (P21-35). Prior to perfusion mice were anaesthetized via intraperitoneal injections of a ketamine (60mg/kg) and xylazine (12mg/kg) cocktail. Once a mouse no longer responded to the pedal reflex, they were transcardially perfused with phosphate-buffered saline (PBS) and then 4% paraformaldehyde (Electron Microscopy Sciences) [PFA] in PBS. Following perfusion, spinal cords were dissected and incubated in 4% PFA for 4h on ice. Spinal cords were then washed in PBS three times for 20mins each on the ice followed by overnight in PBS at 4°C. Subsequently, spinal cords were cryoprotected in 30% sucrose in PBS at 4°C for 2-3 nights. Cryoprotected spinal cords were then embedded in O.C.T compound (Fisher Healthcare) and flash frozen at -55 degrees in a mixture of dry ice and ethanol. Frozen higher lumbar spinal cord segments were sectioned transversely using a cryostat (Leica CM1950) at 30 μm and mounted onto Superfrost Plus Microscope Slides (Fisherbrand).

### Immunohistochemistry

Spinal cord sections were first incubated in PBS containing 0.1% Triton X (PBS-T) for 3 consecutive washes of 5 minutes each. Subsequently, spinal cord sections were incubated in 0.1% PBS-T solution containing primary antibodies and 10% heat-inactivated horse serum (Invitrogen). The primary antibodies used in this study are listed in the Table 1. Following primary antibody incubation, spinal cord sections were washed with PBS for 15 mins (3×5min fresh solution). Sections were then incubated in PBS solution containing secondary antibodies for 1 hour at 4°C. The secondary antibodies used are listed in Table 1. In the end, sections were washed with PBS for 15 mins (3×5min fresh solution) and cover-slipped with fluorescent mounting medium (Dako).

### Retrograde tract tracing of motoneurons

Cholera Toxin Subunit B (CTB) conjugated with Alexa Fluor 488 (0.5mg/1ml PBS) (Molecular Probes) was injected into either proximal iliopsoas (IP), bicep femoris (BF), semitendinosus (ST), tibialis anterior (TA) or gastrocnemius (GS) muscles in P14 *Sim1^Cre+/-^; Ai34^(RCL-Syp/tdT)-^* mice. 7 days after CTB injections, animals were euthanized, and spinal cords were dissected at P21 for further histology study.

### Imaging and reconstruction of V3 synapses onto premotor neurons and motoneurons

Images of retrograde-labelled CTB motoneurons in Sim1^Cre+/-^; Ai34^(RCL-Syp/tdT)-D^ mice were obtained using a Zeiss LSM 510 upright laser scanning confocal microscope with a Zeiss Plan-Apochromat ×63 × 1.4 numerical aperture (NA) objective lens utilizing the tiling and Z-stack function of Zeiss Zen Pro imaging software. The Z-stack images were then imported into Imaris 8.1.2 software for 3D reconstruction and quantitative analysis. The Imaris “Surface” tool was used to reconstruct both the labelled motoneurons (somas and proximal dendrites) and synaptic boutons (Vglut2 terminals and Sim1+ cell terminals). The Imaris “Filter” tool was used to segregate synaptic contacts to only include those which made direct contact with the neuron surfaces. To identify double-labelled TdTom+/VGLUT2+ contacts, the “Find surfaces close to surfaces” MATLAB extension was run in the Imaris software, which allowed for the quantification of colocalized synaptic contacts. TdTom+ and VGLUT2+ boutons were defined as colocalized if the center points of their volumetric reconstructions were within 0.2 microns of each other.

### Optogenetic stimulation of V3 neurons in vitro for neonatal mice

All experiments were performed with postnatal spinal cords from P2-P3 V3ChR2 mice. The mice were anaesthetized, and the spinal cords caudal to thoracic (T) 8 segments were dissected in an ice-cold oxygenated Ringer’s solution (111 mm NaCl, 3.08 mm KCl, 11 mm glucose, 25 mm NaHCO3, 1.25 mm MgSO4, 2.52 mm CaCl2, and 1.18 mm KH2PO4, pH 7.4). The spinal cord was then transferred to the recording chamber to recover at room temperature for 1 h before recording in Ringer’s solution.

Electroneurogram (ENG) recordings of the right (lumbar) L2 and L5 ventral roots, and left L2 – L5 ventral roots made with glass suction electrodes connected via silver-silver-chloride wires to differential amplifier (A-M system, model 1700) with the band-pass filter between 300 Hz and 1 kHz. Analog signals were transferred and recorded through the Digidata 1400A board (Molecular Devices) under the control of pCLAMP10.3 (Molecular Devices). To activate ChR2 in V3 neurons, 488 nm light was delivered by the Colibri.2 illumination system (Zeiss) through 20 × 1.0 numerical aperture (NA) objectives mounted on an up-right microscope (Examiner, Zeiss) onto the ventral surface of the isolated spinal cord. This allowed us to have relatively fixed focal areas. As shown in our previous study^39^, to perform unilateral stimulation, we manually adjusted the field diaphragm and LED light intensity to cover the whole half side of the spinal cord to be the largest illuminated area, and then reduce the illuminated region between approximately one-third to a half of the spinal cord to regulate the number of V3 neurons being activated. After the objective was positioned, a train of 10 X 100ms light pulses was applied with 5 s intervals. The light intensity to produce a minimal ventral root response in V3ChR2 mice was measured and considered the threshold (T). Experiments were conducted with the light at 4xT.

### Window implantation and EMG electrode implantation

To get the light access to the spinal cord, we adopt the window implantation technique ^61,62^. Briefly, mice were anaesthetized (4% isoflurane induction and 2% maintenance, in carbogen; 95% oxygen, 5% CO2) and shaved over the mid-thoracic vertebrae. The surgical area over the thoracic spinal cord was shaved and disinfected with iodine and 70% ethanol. A ∼1.5cm dorsal midline incision was made over the vertebrate T11 to L1, and the muscles between the spinous and transverse processes were resected using scalpel; the animal’s T11 and L1 vertebrates were suspended from a spinal-fork stereotaxic apparatus; muscles that cover the lateral vertebrates from T12 to T13 were removed; two paper clip wires served as anchoring points and were placed onto T12 and T13 vertebrates; cyanoacrylate was carefully applied to all tissue from the edges of the exposed vertebrae to the edges of the cut skin; dental cement was applied over the cyanoacrylate layer to form a rigid ring around the vertebrae and wires; a laminectomy was carried out on T12 and and part of T13 vertebrae to expose the L2 – L3 lumbar spinal cord without damaging any dura matter; liquid Kwik-Sil was applied on the dura matter; and finally a glass window (coverslip) was placed over the spinal cord and sealed with cyanoacrylate and dental cement. Two magnets were imbedded in the dental cement to later attach LEDs (see below).

Two weeks after the window implantation, mice received an EMG implantation as described by Pearson et al. in 2005 and Akay et al. in 2006 and 2014^63–65^. Mice were anaesthetized (4% isoflurane induction and 2% maintenance, in carbogen; 95% oxygen, 5% CO2) and shaved over the thigh, the back of the leg and neck region. The surgical area over the thoracic spinal cord was shaved and disinfected with iodine and 70% ethanol. Fine EMG recording electrodes were implanted into hind-limb muscles, including the flexors of the hip, knee and ankle, respectively iliopsoas (IP), semitendinosus (ST), and tibial anterior (TA), and the extensors of the hip, knee and angle, respectively anterior bicep femoris (BF), vestus lateralis (VL) and medial gastrocnemius (GS). To minimize tissue damage and mechanical distortion of the muscles, small recording electrodes will be used as described in Pearson et al. in 2005^63^. These electrodes cause little interference with leg movements during walking. The wires were led subcutaneously to a small socket embedded in an acrylic headpiece. The headpiece was then stitched to the skin near the neck incision by 5.0 suture. Small incisions were made on the shaved areas and each bipolar electrode was led under the skin from the neck incision to the leg incisions. All cables were bundled in a polyethylene tube (INTRAMEDIC, O.D. 1.57mm). The tube was placed lateral to the spine. Needles on the distal end of each electrode wire were used to draw the pair of electrode wires through the muscle until a knot proximal to a short bared region of the wire was positioned firmly against the surface of the muscle. This positioned a bared wire region in the muscle. The distal end of the pair of electrodes with the needle was knotted at the muscle surface. The wires distal to this second knot were removed by cutting them close to the knot. Afterwards, the incisions in the legs were closed with 5.0 suture.

After each surgery, postoperative analgesia (buprenorphine, 0.03mg/kg) and saline (1ml) were administered, and mice were separated in heated cages until fully awake and monitored for signs of infection or distress. Mice were allowed a recovery period of at least two weeks before experimental procedures commenced.

### Optogenetic stimulation of V3 neurons in vivo

Mice were placed in a custom-built chamber for acclimatization for at least 20 minutes prior to experimentation to minimize stress and basal motor activity. The chamber was designed to restrict gross movements while allowing for comfortable resting and breathing. Ambient temperature was maintained at 22-24°C to ensure thermal comfort.

#### At rest condition

A blue light LED (Marktech, 450 nm with a low dome lens, PN MTE4600N) was used to activate ChR2-expressing V3 neurons through the optical window, with the LED attached to the window during the trial with a magnet embedded in the LED holder. Stimulation began with identifying the threshold light intensity that evoked detectable muscle activity, determined by gradually increasing light intensity in small increments. Following threshold (T; ∼1 mW/mm^2^) determination, the light intensity was adjusted to evoke the full response capacity, which was ∼10xT. Each stimulation consisted of a 15 ms light pulse, with 10-15 trials recorded for each muscle to ensure reproducibility and statistical reliability. Sometimes, an extra 10 trials with 150ms light pules were recorded. Care was taken to ensure that mice remained motionless during light stimulation to attribute recorded muscle activity solely to V3 neuron activation. The EMG signals were recorded using MultiClamp 700B amplifier (Molecular Devices). Analog signals were filtered at 10 kHz with the Digidata 1400A board (Molecular Devices) under the control of pCLAMP10.3 (Molecular Devices). Muscle activity was recorded in a quiet environment to minimize external noise interference.

#### During locomotion condition

Prior to the locomotor task, V3ChR2 mice were acclimatized to walk on custom-made treadmills. No training was performed before any locomotion tests. During the locomotion tests, the mice were placed on the treadmill set to a constant speed at speeds of 15cm/s that elicited consistent voluntary stepping without inducing stress or fatigue. To avoid their fatigue, animals would not perform more than two trials during each experiment, and each trial was <20 s. There was a >1 min rest period between consecutive trials. Once steady locomotion was achieved, as evidenced by consecutive stepping, light stimulation was applied to activate ChR2-expressing V3 neurons. The stimulation protocol consisted of 200 ms light pulses at a frequency of 4 Hz for 5 seconds. The light intensity was adjusted to the same level used in the resting experiment to ensure comparability of the results.

### Ex vivo recording from motoneurons in whole adult spinal cords

We recorded the optogenetically-evoked EPSPs from motoneurons following the methods of Hari et al. ^66^. Adult mice were anaesthetized with urethane (for mice, 0.11 g per 100 g, with a maximum dose of 0.065 g); a laminectomy was performed; and then the entire sacrocaudal spinal cord was rapidly removed and immersed in oxygenated modified artificial cerebrospinal fluid (mACSF composed 118 mM NaCl, 24 mM NaHCO3, 1.5 mM CaCl2, 3 KCl mM, 5 mM MgCl2, 1.4 mM NaH2PO4, 1.3 mM MgSO4, 25 mM D-glucose, and 1 mM kynurenic acid, saturated with 95% O2-5% CO2 and maintained at pH 7.4). Spinal roots were removed, except the sacral S3, S4 and caudal Ca1 VRs and DRs on both sides of the cord. After 1.5 hours in the dissection chamber (at 20 °C), the cord was transferred to a recording chamber containing normal ACSF (nACSF composed of 122 NaCl mM, 24 NaHCO3 mM, 2.5 CaCl2 mM, 3 KCl mM, 1 MgCl2 mM, and 12 mM D-glucose, saturated with 95% O2-5% CO2 and maintained at pH 7.4) maintained at 23–32 °C, with a flow rate >3 ml min−1. A 1-hour period in nACSF was given to wash out the residual anaesthetic before recording, at which time the nACSF was recycled in a closed system. The cord was secured onto tissue paper at the bottom of a rubber (SilGuard) chamber by insect pins in connective tissue and cut root fragments. The cord was oriented with its left side upwards with the ventral surface facing the recording wires.

We employed a grease gap method to record the composite intracellular response of many motoneurons at their exit point to the ventral root, where a high impedance seal on the axon reduces extracellular leak currents, allowing the recording to reflect intracellular motoneuron potentials. We mounted the freshly cut ventral roots onto silver-silver chloride wires just above the bath and covered them in grease over about a 6 mm length, with the grease covering the root up the entry point into the spinal cord. Return and ground wires were in the bath and likewise made of silver-silver chloride. This allowed stable recordings of motoneuron EPSPs, though attenuated by about 50% compared to the intracellular recordings of EPSPs by the voltage-divider law (Re/ (Re + Ra) ∼=1/2, with Re ∼= Ra), where Re and Ra are the extracellular leak resistance and intracellular axial resistance, respectively. The recordings were amplified (2,000 times), high-pass filtered at 0.1 Hz to remove drift, low-pass filtered at 10 kHz, and sampled at 30 kHz (Axoscope 8; Axon Instruments/Molecular Devices, Burlingame, CA). The light used for exciting V3 neurons was from a 447 nm D442001FX laser (Laserglow Technologies, Toronto) focused vertically downward on the V3 neurons. Light was passed through a fiber optic cable (MFP_200/220/900-0.22_2m_FC-ZF1.25 and MFP_200/240/3000-0.22_2m_FC-FC, Doric Lenses) and then a half cylindrical prism the length of about two spinal segments (8 mm; 3.9 mm focal length, Thor Labs), which collimated the light into a narrow long beam (200 µm wide and 8 mm long). The light was adjusted to twice the threshold (T) to produce a ventral root response to V3 neuron activation in spinal cords from V3ChR2 mice (10 - 15 ms, 2T, 447nm, 0.7 mW/mm^2^).

### Locomotor task assessments

After two weeks of EMG implantation surgery, all mice were subjected to perform level-walking, inclined-walking on a treadmill and swimming. At least two trials of every task were recorded. The rest interval between two consecutive trials is at least 1 minute. The whole experiment within one day did not exceed two hours of testing spread out over the day. During any given run, if the mouse hits the back wall of the chamber, the treadmill will be stopped immediately. If the animal is resisting running or cannot catch up with the treadmill speed, gentle stimulation, like blow or touch the hind body, will be applied. The EMG signals were recorded using Power1401 (model MA 102, custom-built in the workshop of the Zoological Institute, University of Cologne, DEU) and Spike 2 (Version 7.09a, Cambridge Electronic Design) software.

#### Level and inclined treadmill walking

Mice were placed on a motorized treadmill set to a level surface. The speed was gradually increased to 15 cm/s that each mouse could sustain. Then the treadmill was inclined to a 17.5° angle to increase the locomotor challenge. Mice were observed for their ability to keep up with the treadmill speed, and performance metrics were recorded.

#### Swimming

A clear, custom-made Plexiglass tank (100 x 10 x 40cm) was filled with warm water (23-26℃) to a depth of 25-30cm. A platform was placed at one end so that it was clearly visible to the mouse from the opposite end of the tank. Mice were lowered carefully into the water and allowed to swim to the platform. Each mouse underwent five trials per session, and each trial was limited to 1 min in duration. Between trials, the mice were given a 2-minute rest, during which they were placed in a clean, dry cage with a paper towel on the top of a heating mat set to 37 ℃. Initially, some mice required a short training period during which the distance required for the mouse to reach the platform was increased until the mouse is able to swim the full distance. Mice were carefully monitored during the test. Mice displaying underwater tumbling or any distressing behaviour were scooped up and rescued immediately. For drying the mice following the testing session, the animals were placed in an empty cage with some paper towels on top of a heating mat set to 37 ℃. The Plexiglass tank was disinfected with Peroxigard (Peroxigard™, Virox® Technologies Inc) before and after the group test started and finished.

### Data analysis

#### V3 synapse density on motoneurons

Quantitative synaptic input data was taken from Imaris software. The number of identified double-labelled TdTom+/VGLUT2+ synaptic contacts was then divided by the total number of Vglut2+ synaptic contacts to yield the percentage of VGLUT2+ synapses onto each motoneuron supplied by Sim1+ cells.

#### ENG and EMG data

The recordings were initially rectified and smoothed (τ = 0.05 s) in Spike 2. All processed data were then transformed to MATLAB (Version R2019a, The MathWorks, Inc.). The optical stimulation was recognized as a TTL signal sent from Digidata 1400A board (Molecular Devices) that synchronizes to the laser controller (brand name). To measure the response from the ventral root evoked by V3 activation, we measured the integrated ENG (iENG) 20 ms before light and after the first spike onset during light exposure. To measure the muscle activity evoked by V3 activation, we measured the mean integrated EMG 20ms after light exposure. Considering the difficulties in comparing absolute EMG amplitude between animals, we normalized each EMG to its peak observed during voluntary activity (e.g. walking) to account for inter-animal variability. The equation for this is shown below.

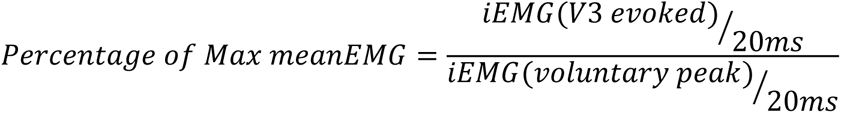

To compare the EMG activity in wildtype and V3OFF animals in different locomotor tasks, we measured iEMG and mean iEMG as described in Figure 6Ci. To identify if the muscle activity we measured was in one step cycle, we manually tracked the toe touching and lifting point of each step in the video recording to decide the EMG activities from different muscles in one step cycle. Then, we manually marked the onset and offset of muscle activity to calculate the burst duration, as shown in Figure 6Ci.

To measure the latency of activities evoked by V3 activation in in vitro and in vivo experiments, signal traces in all trials in one animal were averaged with the reference to the light onset. The time of the first detectable augment was marked. The latency is the time interval between the onset of the light and the onset of this augment.

### Statistics

The statistical analysis was performed in Prism7 (GraphPad Software, Inc.) and MATLAB (Version R2018a, The MathWorks, Inc). One-way ANOVA was used to compare the levels of Sim1+/Vglut2+ inputs onto motoneurons belonging to different motor pools and acting at different joints and Tukey HSD test was used for multiple comparisons.

Unpaired t-test was used to compare between flexor and extensor motor pools. Wilcoxon test and paired t test were used to compare the iENG activity between before and after V3 activation, the latency for nerves and muscles, and the iEMG activity between each muscle. Student t-test and Welch’s t-test were used to compare the differences between WT and V3OFF mice’s EMG burst duration and activities in three locomotor tasks.

## Acknowledgements

The authors are thankful for the generous gifts of Sim1^Cre/+^ mouse from Dr. Martyn Goulding (Salk Institute for Biological Studies, CA, USA). The research was funded by the Canadian Health Institutes of Research (MOP 14697 and PJT 165823 to D.J.B.), (PJT-162357 to TA), and (MOP-110950 and PJT-173547 to Y.Z.), the US National Institutes of Health (R01NS47567 to D.J.B. and K.F. and R01 NS115900 to TA), and the Natural Sciences and Engineering Research Council of Canada (RGPIN 04880 to Y.Z.).

## Contributions

H. Z., D.D.-G, K.K.F., D.J.B and Y.Z. conceptualized the paper; D.D.-G, and C.S.M. designed and performed histology, confocal microscopy, and 3D reconstruction; H. Z. and J.B.-F. designed and performed in vitro whole cord recordings; A.M.L-O. and K.H. designed and performed in vitro motoneuron recordings; H. Z., K.K.F., R.L.L. and D.J.B designed, performed, and analyzed in vivo optogenetic and behavioural experiments; H. Z., D.D.-G and C.S.M. analyzed the data and performed all statistics. H. Z., D.J.B and Y.Z. wrote the paper, with editing from other authors. These authors contributed equally: H. Z., D.D.-G. These authors equally supervised this work as senior authors: Y.Z., K.K.F., T.A. and D.J.B.

