## Supplementary Table for "Widespread innervation of motoneurons by spinal V3 neurons globally amplifies locomotor output in mice"

| Percentage of total Vglut2 contacts | IP (%) | BF (%) | ST (%) | VL (%) | TA (%) | GS (%) |
|---|---|---|---|---|---|---|
| Avg | 25.47 | 23.14 | 12.75 | 26.60 | 21.92 | 16.19 |
| Stdev | 19.24 | 13.20 | 5.80 | 16.23 | 14.37 | 8.36 |
| | Hip (%) | | Knee (%) | | Ankle (%) | |
| Avg | 24.23 | | 19.91 | | 19.42 | |
| Stdev | 18.54 | | 13.06 | | 12.42 | |
| | Flexor (%) | | | Extensor (%) | | |
| Avg | 22.08 | | | 20.73 | | |
| Stdev | 16.10 | | | 14.72 | | |
